# Trans-generational effects on diapause and life-history-traits of an aphid parasitoid

**DOI:** 10.1101/582932

**Authors:** Tougeron K., Devogel M., van Baaren J., Le Lann C., Hance T.

## Abstract

Transgenerational effects act on a wide range of insects’ life-history traits and can be involved in the control of developmental plasticity, such as diapause expression. Decrease in or total loss of winter diapause expression recently observed in some species could arise from inhibiting maternal effects. In this study, we explored transgenerational effects on diapause expression and traits in one industrial and one wild strain of the aphid parasitoid *Aphidius ervi*. These strains were reared under short photoperiod (8:16 h LD) and low temperature (14 °C) conditions over two generations. Diapause levels, developmental times, physiological and morphological traits were measured. Diapause levels increased after one generation in the wild but not in the industrial strain. For both strains, the second generation took longer to develop than the first one. Tibia length and wing surface decreased over generations while fat content increased. A crossed-generations experiment focusing on the industrial parasitoid strain showed that offspring from mothers reared at 14 °C took longer to develop, were heavier, taller with wider wings and with more fat reserves than those from mothers reared at 20 °C (8:16 h LD). No effect of the mother rearing conditions was shown on diapause expression. Additionally to direct plasticity of the offspring, results suggest transgenerational plasticity effects on diapause expression, development time, and on the values of life-history traits. We demonstrated that populations showing low diapause levels may recover higher levels through transgenerational plasticity in response to diapause-induction cues, provided that environmental conditions are reaching the induction-thresholds specific to each population. Transgenerational plasticity is thus important to consider when evaluating how insects adapt to changing environments.

## Introduction

Transgenerational plasticity (TGP) represents a particular type of phenotypic plasticity meaning that environments experienced by the grand-maternal or maternal generation generate phenotypic variations in the offspring generations (Mousseau and Dingle, 1991). TGP effects on insect ecology and life history traits are quite common and have been proved to be adaptive when the parental environment accurately predicts or is similar to the offspring’s environment (Sgrò et al., 2016). Among other mechanisms, TGP can involve epigenetics (Youngson and Whitelaw, 2008), which is known to act on phenotypic expression and reaction norms over several generations (Reynolds, 2017; Uller, 2008).

Maternal effects have been particularly well studied in the context of developmental plasticity in insects (Mousseau and Fox, 1998; Uller, 2008). In insects, facultative diapause, a phase of developmental arrest and low metabolic activity, is induced mostly by environmental conditions encountered by the overwintering generation (Tauber et al., 1986). Experienced environmental cues underwent by the mother females also inform for deleterious conditions to come. During ovogenesis, these conditions influence carbohydrates and polyols contents of the egg which acts on the developmental plasticity of the embryo (Denlinger, 2002). In this case the sensitive stage is the ovipositing female and diapause is initiated in its offspring (Saunders, 1965; Lacour et al., 2015; Tougeron et al., 2017a). In most cases, both maternal and diapausing generation plasticity are involved in determining diapause incidence (Tauber et al., 1986).

Powell & Bale (2008) and Coleman *et al.* (2014) demonstrated the existence of adaptive maternal effects on cold hardening in overwintering insects, respectively in the grain aphid *Sitobion avenae* (Hemiptera: Aphididae) and the blow fly *Calliphora vicina* (Diptera: Calliphoridae). Among parasitoid insects, Voinovich et al., (2015) showed that diapause was induced by maternal effects in *Trichogramma* (Hymenoptera: Trichogrammatidae) while it was averted in the offspring when mothers were reared at high temperatures. In some species, there is a non-photo-thermo-sensible generation at which it is impossible to induce diapause, due to maternal effects. This is a potential adaptation to avoid diapause induction in spring during which environmental conditions are similar to fall (Reznik and Samartsev, 2015).

Phenotypic plasticity across generations is an important mechanism for organisms to cope with climate variations at different temporal scales. In the context of rapid climate change and increase in temperature fluctuation and unpredictability, a better knowledge about response to environmental variations through TGP may help us to better understand the ability of rapid adaptation of key species in ecosystems (Donelson et al., 2018). There is growing evidences that, in response to climate warming, or when reared in the laboratory for a long time at constant favorable temperature conditions, insects can reduce or abort winter diapause (Musolin, 2007; Tougeron et al., 2017b). For instance, the parasitoid *Binodoxys communis* (Hymenoptera: Braconidae) lost its capacity to enter diapause in less than 300 generations maintained in the laboratory (Gariepy et al., 2015). These shifts in overwintering strategies can be triggered either by (i) evolutionary (genetic) changes (e.g., Bradshaw & Holzapfel 2001) on diapause induction thresholds or (ii) by plastic responses, when insects receive an improper environmental signal to enter diapause, which could include developmental TGP and bet-hedging strategy (Hopper, 1999; Tougeron et al., 2018a).

*Aphidius* parasitoid wasps are interesting models to study adaptive changes in diapause expression. In this genus, some maternal effects on diapause induction have been reported, confirming that both parental and direct induction can occur (Brodeur and McNeil, 1989; Langer and Hance, 2000; Tougeron et al., 2017a). Recent studies indicated a decrease in diapause expression in some *Aphidius* populations, in both natural and laboratory conditions (Tougeron et al., 2017b). We questioned whether a low proportion of diapausing individuals in some *Aphidius* populations could be due to a genetic loss of diapause or to the inhibition of plastic diapause expression due to the maternal environment. The first aim of this study was to measure the consequences of maternal effects on diapause incidence, developmental time and life-history traits, as parasitoid mothers could induce or inhibit diapause in their offspring and affect their traits depending on the environmental signals they perceive. The second aim was to investigate the propensity of parasitoids to increase diapause levels and modify trait value over several generations exposed to diapause-inductive conditions.

To achieve these goals, we chose two contrasted strains of the parasitic wasp *Aphidius ervi* Haliday (Hymenoptera: Braconidae) regarding diapause induction. The first one has been reared for mass rearing production by a biological control company under non-diapausing conditions for decades. It expresses barely no diapause (<5%) (Mubashir-Saeed *et al.* in prep) when exposed to short photoperiod and low temperature conditions. The second strain is a Canadian wild strain known to express high diapause levels (>90%) (Tougeron et al., 2018a; Tougeron et al., 2018b). Two main experiments were realized: (1) we exposed these two strains to known diapause-inducing conditions during two consecutive generations. (2) We exposed the maternal generation of the industrial strain to either diapause-inductive or non-diapause-inductive conditions to test for maternal effects on the offspring. We predict that if diapause plasticity is conserved, diapause levels (1) should increase across generations exposed to diapause-inductive conditions and (2) when mothers are exposed to diapause-inductive conditions. We also predict that life-history traits and development times of individuals not only depend on the conditions they experience but also on the thermal experience of the precedent generation.

## Material and methods

### Insect production

Rearing cultures were started with two strains of *A. ervi*: an industrial one provided by the Viridaxis SA company (Charleroi, Belgium), and a wild Canadian strain collected in the pea and clover fields in summer 2015 (Québec, 45.584 °N, 73.243 °W). It is important to note that *industrial* and *wild* are used as generic terms to indicate that the first one has been maintained for significantly more generations under standard rearing conditions than the second, which is the factor that actually differentiates both strains and from which an effect is expected. Both strains were maintained on parthenogenetic pea aphids *Acyrthosiphon pisum* Harris (Hemiptera: Aphididae) provided by Viridaxis SA and reared on faba bean (*Vicia faba*: Fabaceae) in pots of potting soil treated with diflubenzuron 25%, an insecticide for shore flies. Insects and plants were maintained at 20 °C, 75 ± 5% Relative Humidity (RH) and with a long day photoregime of 16:8 h Light:Dark (LD).

Mummies (i.e., dead aphid containing a parasitoid prepupa) were collected from the rearing culture and isolated in Eppendorf tubes (1.5 mL) until parasitoid emergence. Within the first 48 h after emergence, batches of five female and five male parasitoids (Generation 0, G0) were placed in Petri dishes (15 cm diameter) for 24 h for mating, with access to honey and water for feeding. To produce the first generation (G1), female parasitoids of each batch were then allowed to parasitize a cohort of 100 (± 10) parthenogenetic *A. pisum* second instar aphids on a pot of *V. faba* bean, closed by a piece of fine netting, and with an access to honey and water, for 48 h. Mating and oviposition of G0 were both conducted under standard rearing conditions. Then, the pots with parasitized aphids were placed in climate chambers for the experiments (see experimental design for the different treatments) and, once formed, mummies were isolated in Eppendorf tubes (1.5 mL) until adult emergence (G1). A part of these newly emerged G1 adult was freeze-killed for further trait measurements and the other part was used to produce the second generation (G2) as explained above for the first generation and according to the different thermal treatments (see below).

### Experimental design

#### Diapause and life-history traits over generations

Individuals of both parasitoid strains were reared for two successive generations at 14 °C and 8:16 h LD photo-regime during their entire development. A third generation was produced under the same conditions for the industrial strain to confirm the trend observed over two generations. The selected conditions are known to induce diapause in Aphidiinae parasitoids from temperate areas (Christiansen-Weniger and Hardie, 1999; Tougeron et al., 2017b; Sigsgaard, 2000). This experiment was replicated in three climate chambers to which pots of parasitized aphids were randomly assigned at each generation.

#### Maternal effects

In parallel, maternal effect was tested using the industrial strain only. For this purpose, parasitoids from the rearing culture were randomly assigned to different thermal conditions during their entire development (G1: 14 °C or 20 °C at 8:16 h LD) and their offspring (G2, within the parasitized aphids) were placed at 14 °C, 8:16 h LD to have lineages of different temperature history. The temperature of 20 °C does not induce diapause when associated with short photoperiod regime (Tougeron et al., 2017b) and it is different enough from 14 °C to expect differences in traits, development time and diapause incidence through maternal effects in *A. ervi* (Malina and Praslicka, 2008). Three climate chambers were used to randomly assign the aphid pots.

### Measures

#### Emergence patterns and diapause

For each replicate of a same cohort (i.e., individuals of a same generation, strain and temperature treatment), we measured time from oviposition to first emergence. We also measured the sex ratio in emerged individuals and the proportions of emerged, dead and diapausing individuals. Emergences were checked once a day at the same hour and within 24h after emergence adult parasitoids were frozen in liquid nitrogen and stored at −80 °C for further analyses. Mummies from which no parasitoid emerged 10 days after the last adult emergence in the same cohort were dissected to verify if they were in diapause (gold-yellow plump parasitoid larva) or dead (after the protocol of Tougeron et al., 2017b).

#### Fitness-related traits

We analyzed morphological trait indicators of fitness in parasitoids species on individuals of each cohort and treatment (generation, strain, or maternal origin). Traits were measured on an equal number of females and males. The length of the front right tibia, an estimation of adult size (Ellers, 1995) and the surface of the right wing were measured by image analysis method. Digital pictures of tibia and wing were captured with a camera (Sony N50) mounted on a stereomicroscope and pictures were analyzed using the numeric image analysis software ImageJ (Wayne Rasband, USA). Individuals were also weighted with an electronic precision balance (Mettler-Me22; sensitivity: 1µg) to obtain fresh mass. Their dry mass (DM) was measured after they were dried out at 60 °C during 3 days in oven. Each dried individual was placed in an Eppendorf tube (1.5 mL) containing 1 mL of a chloroform:methanol solution (1:2) and left for two weeks on agitator. They were dried out again for 12 h in oven at 60 °C to eliminate the residues of the solution before being weighted once again to obtain the lean dry mass (LDM). Fat content (FC), a relevant measure of survival (Ellers, 1995), was then obtained using the formula : Fat mass (FM) = DM–LDM and FC= FM/LDM (Colinet et al., 2007b).

### Statistical analyses

All statistical analyses were carried out using the R software (R Development Core Team 2017). Differences in emergence patterns and morphological traits between cohorts were assessed by testing the effect of the factors “strain”, “generation”, their interaction and the factor “sex” in the experiment on transgenerational effects. Data of the third generation of the industrial strain was compared to the first two generation of the same strain using separate models. The effect of the explanatory variable “temperature history” was tested in the maternal effect experiment (i.e., combination of maternal treatment origin and experienced temperature by the offspring). Emergence patterns were fitted to a survival logistic-rank model (using package *survival*) by adding a censoring factor for non-emerged living mummies and models were compared using a likelihood-ratio chisquare method. Proportions of emerged, dead and diapausing individuals were compared between cohorts by fitting generalized linear models based on a quasibinomial distribution and a logit link function for each parameter, and models were compared using a Fisher statistic. Data of each morphological trait (fresh mass, tibia length, wing surface and fat contents) was fitted to linear mixed models (package *nlme*), and the climate chambers identity was considered as a random factor in these models. The significance of each explanatory variable was tested using the ‘Anova’ function (package *car*, type II ANOVA). Analyses were followed by Tukey multiple comparison post-hoc tests (using package *multcomp* and *lsmeans*), in order to assess differences among modalities of significant variables.

## Results

### Diapause and life-history traits over successive generations

#### Emergence patterns and diapause

Patterns of emergence (at 14 °C, 8:16 h LD) were significantly different between generations, with the second generation taking longer to develop than the first one (Survival, LR=52.97, p<0.001, mean emergence time of 6.23 ± 0.16 and 8.42 ± 0.31 days after first emergence, respectively). There were no difference between strains (Survival, LR=2.58, p=0.108), which both exhibited the same variation between generations (Survival, LR=0.26, p=0.608). Parasitoids from the third generation of the industrial strain took an average of 11.28 ± 0.49 days to emerge, after first day of parasitoid emergence, which is longer than the first (TukeyHSD, z=11.1, p<0.001) and second generation (TukeyHSD, z=5.1, p<0.001) (Figure 1). In the wild strain it took an average of 27.33 ± 1.85 (mean ± SE) days in the first generation and 27.00 ± 1.15 days in the second to get the first emergence after oviposition. For the industrial strain, it took an average of 22.67 ± 0.67 days before the first emergences in the first generation, 27.33 ± 1.33 days in the second and 27.67 ± 0.33 days in the third.

**Figure 1:**
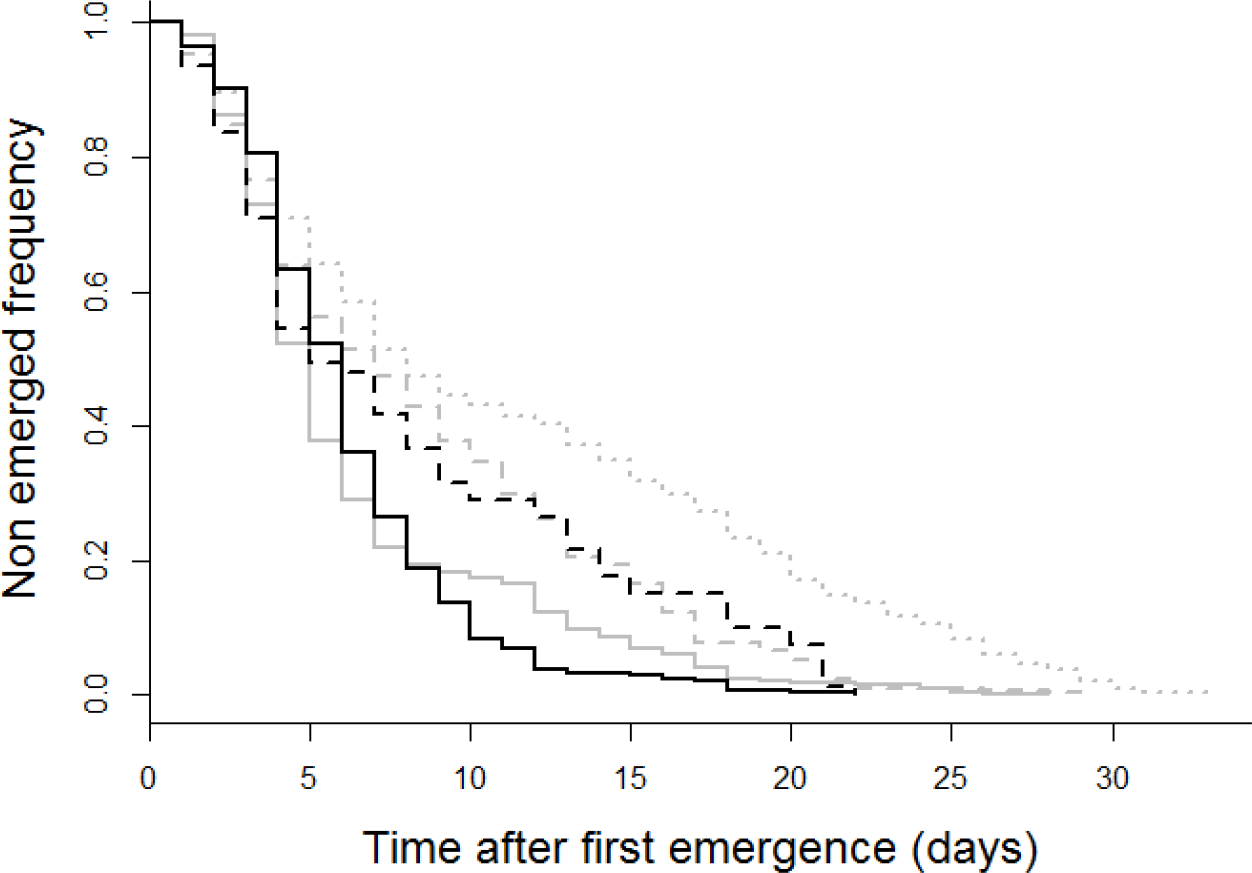
Emergence patterns from first parasitoid emergence (at day 1) in each strain, wild (black) and industrial ones (grey), in the 1^st^ (full lines N=283 and N=426 respectively), 2^nd^ (dashed lines, N=79 and N=302 respectively), and 3^rd^ (point line, N=298, industrial strain only) generations.

The proportion of parasitoids entering diapause was significantly different between strains (GLM, F=6.16, p<0.05) and generations (GLM, F=9.59, p<0.05). Levels of diapause were overall lower in the industrial strain (TukeyHSD, z=−1.96, p<0.05), and in the first generation (TukeyHSD, z=2.96, p<0.005). There was an interaction effect between strain and generation (GLM, F=6.95, p<0.05). Levels of diapause statistically increased in the wild strain (Tukey, z=2.96, p<0.05), but were not different in the industrial one (TukeyHSD, p=0.99, z=0.26) between the first and second generations (Table 1). No parasitoid from the third generation of the industrial strain entered diapause.

**Table 1:** means ± SE of sex ratio and percentage of emerged, dead and diapausing individuals for the first and second generation of both strains and the third generation of the industrial strain (W: Wild strain and I: Industrial strain.

| Generation | Strain | N | Sex ratio | % Emerged | % Dead | % Diapause |
| --- | --- | --- | --- | --- | --- | --- |
| 1 | W | 476 | $0.65 \pm 0.04$ | $52.73 \pm 21.33$ | $38.30 \pm 17.20$ | $8.97 \pm 4.45$ |
| | I | 615 | $0.14 \pm 0.02$ | $69.21 \pm 5.42$ | $30.00 \pm 5.88$ | $0.79 \pm 0.46$ |
| 2 | W | 291 | $0.57 \pm 0.27$ | $22.47 \pm 4.47$ | $44.74 \pm 5.50$ | $21.66 \pm 9.97$ |
| | I | 422 | $0.52 \pm 0.09$ | $69.35 \pm 6.77$ | $30.05 \pm 6.43$ | $0.60 \pm 0.60$ |
| 3 | I | 424 | $0.35 \pm 0.05$ | $67.00 \pm 7.22$ | $33.00 \pm 7.22$ | 0.00 |

The same proportion of individuals emerged in both strains (GLM, F=4.11, p=0.07) and generations (GLM, F=0.93, p=0.36), and the same proportion died in both strains (GLM, F=0.67, p=0.44) and generations (GLM, F=0.14, p=0.72) (Table 1). There were significant differences of sex ratio between strains (GLM, F=14.6, p<0.01) and a significant interaction between strains and generations (GLM, F=7.62, p<0.05). There was a higher proportion of males in the wild strain than in the industrial one (Tukey, z=4.49, p<0.001) at the first generation, but both strains presented a balanced sex-ratio at the second generation (TukeyHSD, z=0.20, p=0.99) (Table 1).

#### Fitness-related traits

There were significant morphological differences between sexes for both strains and generations, females being heavier than males (LMM, χ^2^=19.8, p<0.001, mean ± SE 0.43 ± 0.01 mg *vs.* 0.33 ± 0.01 mg), having longer tibias (LMM, χ^2^=52.5, p<0.001, 0.53 ± 0.01 mm *vs.* 0.49 ± 0.01 mm) and a higher fat content ratio (χ^2^=15.5, p<0.001, 0.36 ± 0.02 *vs.* 0.24 ± 0.02). There was no difference in wing surfaces between sexes for both strains and generations (LMM, χ^2^=0.22, p=0.64, 1.14 ± 0.01 mm^2^ *vs.* 1.16 ± 0.02 mm^2^). The same trend is observed for the third generation of the industrial strain: fresh weight 0.41 ± 0.01 mg vs. 0.33 ± 0.01 mg, tibia length 0.50 ± 0.01 mm vs. 0.45 ± 0.01 mm, fat content ratio 0.29 ± 0.02 vs. 0.19 ± 0.02, wing area 1.26 ± 0.02 mm^2^ vs. 1.25 ± 0.02 mm^2^, for females and males, respectively.

Parasitoids of the wild strain had overall lower fresh weights than industrial ones (LMM, χ^2^=73.29, p<0.001, 0.32 ± 0.01 mg *vs.* 0.43 ± 0.01 mg, respectively) and this difference remained the same at each generation (LMM, χ^2^=0.02, p=0.88). There were no differences in fresh weights between generations, all strain confounded (LMM, χ^2^=3.38, p=0.07, 0.43 ± 0.01 mg *vs.* 0.34 ± 0.01 mg, for G1 and G2, respectively). The third generation of the industrial strain had a mean fresh weight of 0.34 ± 0.01 mg, which was lower than G1 (TukeyHSD, z=−8.9, p<0.01) but similar to G2 (TukeyHSD, z=− 0.9, p=0.61) (Figure 2A)

**Figure 2:**
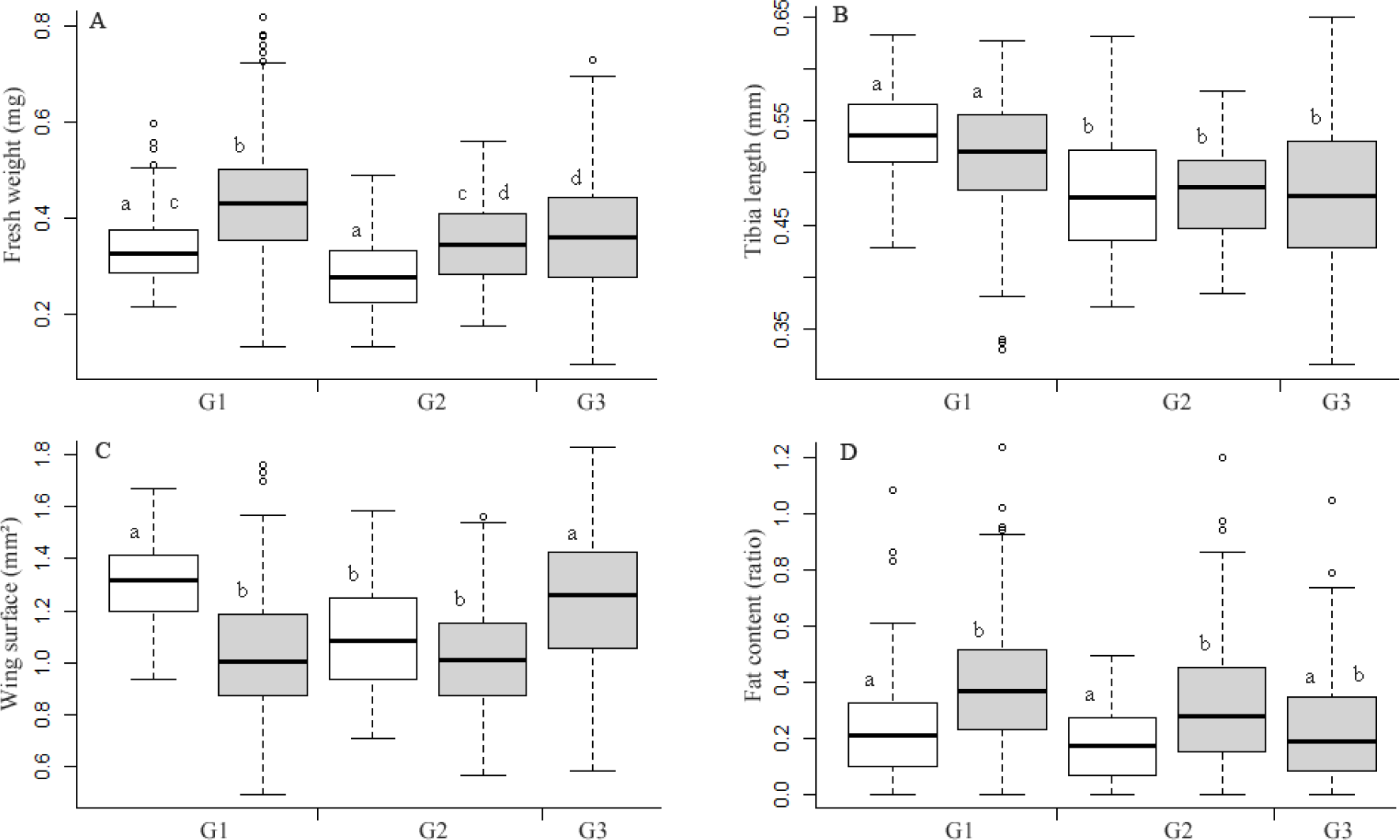
Mean trait values of individuals (± CI 95 %). **A**: fresh weight (mg), **B**: tibia length (mm), **C**: wing surface (mm^2^) and **D**: fat contents (ratio fat mass/lean dry mass) in wild (white) and industrial (grey) strains of *Aphidius ervi* reared at 14 °C, 8:16 h LD for the 1^st^ generation (G1, left), 2^nd^ generation (G2, middle) and 3^rd^ (G3, right, for the industrial strain only). N=81, 151, 70, 105, 142 from left to right, respectively. Lowercase letters indicate significant differences (TukeyHSD post-hoc tests).

There was no significant difference of tibia length between both strains (LMM, χ^2^=0.33, p=0.56), and parasitoids had overall smaller tibias at the second generation (LMM, χ^2^=4.71, p<0.05, 0.54 ± 0.01 mm *vs.* 0.49 ± 0.01 mm, for G1 and G2, respectively). The third generation of the industrial strain had a mean tibia length of 0.48 ± 0.01 mm, which was lower than G1 (TukeyHSD, z=−8.1, p<0.01) but similar to G2 (TukeyHSD, z=−1.1, p=0.53) (Figure 2B)

There were significant differences of wing surface between both strains (LMM, χ^2^=35.9, p<0.001) and generations (LMM, χ^2^=31.3, p<0.001), with a significant interaction (LMM, χ^2^=9.84, p<0.001). Wild strain individuals had overall wider wings than industrial ones (1.23 ± 0.02 mm^2^ *vs.* 1.10 ± 0.01 mm^2^, respectively). Wing surfaces of wild strain individuals decreased at the second generation (TukeyHSD, t.ratio=5.60, p<0.001) while those of the industrial strain remained the same (TukeyHSD, t.ratio=2.44, p=0.07). The third generation of the industrial strain had a mean wing surface of 1.16 ± 0.02 mm^2^, which was higher than G1 (TukeyHSD, z=4.6, p<0.01) and G2 (TukeyHSD, z=6.5, p<0.001) (Figure 2C).

Parasitoids from the wild strain had lower fat contents ratio than parasitoids from the industrial strain (LMM, χ^2^=8.12, p<0.001, ratio of 0.22 ± 0.02 *vs.* 0.37 ± 0.01, respectively). There was no difference between generations (LMM, χ^2^=2.0, p=0.15). The third generation of the industrial strain had a mean fat content ratio of 0.25 ± 0.02, which was similar to G1 (TukeyHSD, z=−2.7, p=0.07) and G2 (TukeyHSD, z=−2.2, p=0.08) (Figure 2D).

### Maternal effect

#### Emergence patterns and diapause

This experiment was done on the industrial strain only. Patterns of emergence were significantly different between temperature conditions and generations (Survival, LR=495.34, p<0.001) (Figure 3). For the maternal generation (G1), it took an average of 15.7 ± 1.8 (mean ± SE) days after eggs from G0 were laid to get the first emergences at 20 °C, while it took 22.67 ± 0.67 days at 14 °C. In the offspring generation (G2) exposed to 14°C, parasitoids from mothers (G1) reared at 14 °C had a longer mean development time than those from mothers (G1) reared at 20 °C (8.51 ± 0.35 days *vs.* 5.03 ± 0.39 days, respectively, from first day of parasitoid emergence). First emergences were also observed later in individuals from mothers reared at 14 °C than from mothers reared at 20 °C (27.33 ± 1.30 days *vs.* 26.00 ± 1.10 days respectively) (Figure 3).

**Figure 3:**
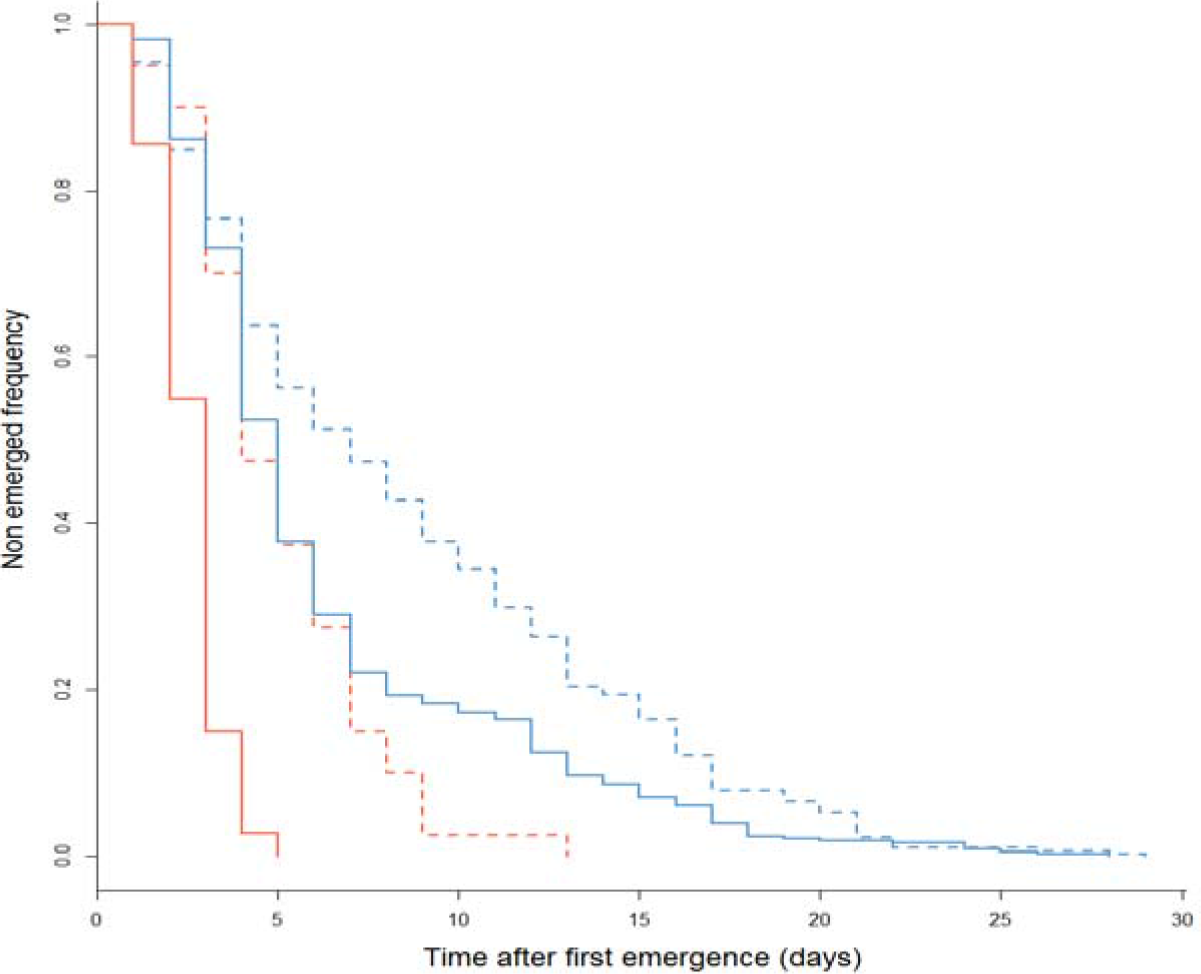
Emergence patterns from first parasitoid emergence (day 1) of maternal generations reared at 14 °C (full blue line, N=426) or 20 °C (full red line, N=180) and offspring reared at 14 °C from either mothers reared at 14 °C (dashed blue line, N=302) or 20 °C (dashed red line, N=40) of an industrial strain of *Aphidius ervi*.

Among cohorts (i.e., parasitoid lineage history), similar proportions of individuals emerged (GLM, F=1.67, p=0.26), died (GLM, F=0.47, p=0.30) and entered in diapause (GLM, F=0.97, p=0.46) (Table 2). Sex ratios differed between conditions (GLM, F=10.53, p<0.01), as mothers reared at 14 °C produced more males than mothers reared at 20 °C (Table 2).

**Table 2:** Means ± SE of sex ratio and percentage of emerged, dead and diapausing individuals for each generation of the industrial strain of *Aphidius ervi*. Data show trait measurements on parasitoids of the first (maternal) generation reared either at 14 °C or 20 °C and of the second (offspring) generation reared at 14 °C but produced by mothers from 14 °C or 20 °C. The photoregime was 8:16 h LD for each temperature condition.

| Mothers' rearing temperature | Offspring Rearing temperature | N | % Emerged | % Dead | % Diapause | Sex-ratio |
| --- | --- | --- | --- | --- | --- | --- |
| G0: 20°C | G1: 14 °C | 615 | 69.21 ± 5.42 | 30.00 ± 5.88 | 0.79 ± 0.46 | 0.14 ± 0.02 |
| G0: 20°C | G1: 20 °C | 212 | 83.97 ± 3.92 | 16.03 ± 3.92 | 0.00 | 0.21 ± 0.08 |
| G1: 14 °C | G2: 14 °C | 422 | 69.35 ± 6.77 | 30.05 ± 6.43 | 0.60 ± 0.60 | 0.52 ± 0.09 |
| G1: 20 °C | G2: 14 °C | 71 | 69.66 ± 9.93 | 49.43 ± 10.67 | 0.91 ± 0.74 | 0.23 ± 0.05 |

#### Fitness-related traits

Globally, there were differences of fresh weight (LMM, F=27.03, p<0.001) (Figure 4A), tibia length (LMM, F=19.39, p<0.001) (Figure 4B), wing surface (LMM, F=25.31, p<0.001) (Figure 4C), and a marginally significant difference in fat content (LMM, p=0.05, F=7.64) (Figure 4D) between maternal cohorts and between offspring female cohorts (parasitoid lineage history).

**Figure 4:**
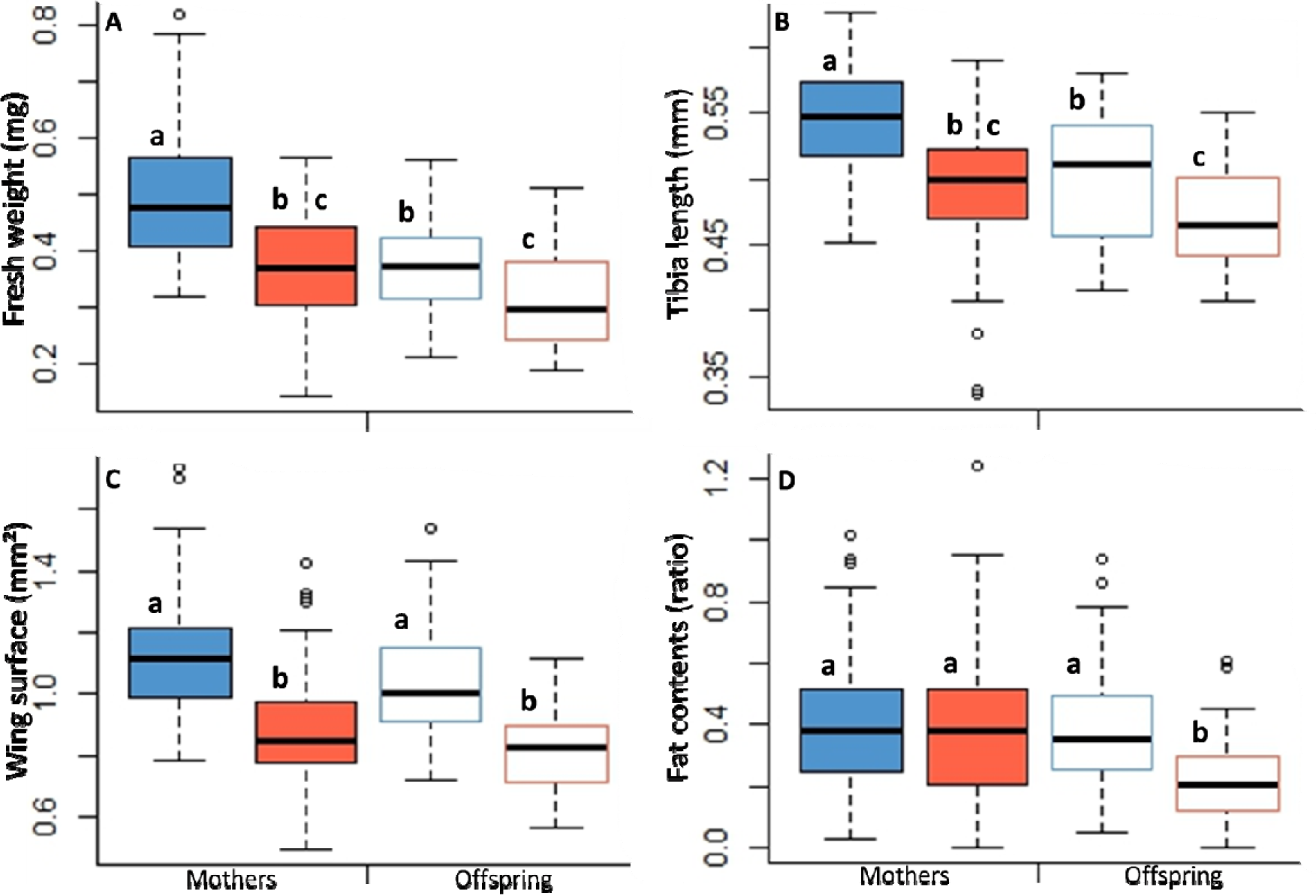
Mean trait values of individuals (± CI 95 %). **A**: fresh weight (mg), **B**: tibia length (mm), **C**: wing surface (mm^2^) and **D**: fat contents (ratio fat mass/lean dry mass) for the different thermal conditions and generations of an industrial strain of *Aphidius ervi*. Left, the maternal generation reared at 14°C (filled in blue) (N=151) and 20°C (filled in red) (N=100). Right, the second generation (offspring) reared at 14°C but from mothers reared at 14°C (blue line) (N=105) or 20°C (red line) (N=31). Lowercase letters indicate significant differences (TukeyHSD post-hoc tests).

Mothers reared at 14 °C were significantly heavier than mothers reared at 20 °C (TukeyHSD, z=3.16, p<0.01, 0.49 ± 0.01 mg *vs.* 0.35 ± 0.01 mg, respectively). Offspring of mothers reared at 14 °C were heavier than those of the mothers reared at 20 °C (TukeyHSD, z=3.02, p<0.05, 0.36 ± 0.01 mg *vs.* 0.29 ± 0.02 mg) (Figure 4A).

Mothers reared at 14 °C had longer tibias than mothers reared at 20 °C (TukeyHSD, z=3.13, p<0.01, 0.54 ± 0.01 mm *vs.* 0.49 ± 0.01 mm, respectively), and their offspring were bigger than those from mothers reared at 20 °C (TukeyHSD, z=3.14, p<0.05, 0.50 ± 0.01 *vs.* 0.46 ± 0.01, respectively) (Figure 4B).

Mothers reared at 14 °C had significantly wider wing surface than mothers reared at 20 °C (TukeyHSD, z=6.52, p<0.001, 1.13 ± 0.02 mm^2^ *vs.* 0.89 ± 0.02 mm^2^, respectively), and their offspring had wider wing surface than those from mothers reared at 20 °C (TukeyHSD, z=4.81, p<0.001, 1.07 ± 0.02 mm^2^ *vs.* 0.83 ± 0.03 mm^2^, respectively) (Figure 4C).

Mothers reared at 14 °C had a similar fat content ratio than mothers reared at 20 °C (TukeyHSD, z=0.58, p=0.9, 0.38 ± 0.02 *vs.* 0.37 ± 0.03, respectively), and their offspring had a marginally significantly higher fat content ratio than those from mothers reared at 20 °C (TukeyHSD, z=0.10, p=0.05, 0.36 ± 0.02 *vs.* 0.21 ± 0.03, respectively) (Figure 4D).

## Discussion

We predicted that diapause levels should increase when parasitoids are exposed to diapause-inductive conditions over generations. Diapause levels increased over two generations in the Canadian wild parasitoid strain (although diapause levels were overall low), but not in the industrial strain, even after three generations. Results on the wild strain showed that higher diapause levels can be retrieved after only one generation, provided that induction thresholds are sensitive enough to environmental cue. Both transgenerational plasticity and direct induction plasticity are involved in diapause induction in the wild strain of *A. ervi*. Results on the industrial strain suggest either that most of the parasitoids lost their capacity to enter diapause as an adaptation to standard rearing conditions, or that conditions did not reach the threshold necessary to trigger developmental plasticity in this population. Both strains exhibited a lengthening of the emergence time and changes in fitness-related traits over the two generations, as hypothesized. The observed trend for the third generation in the industrial strain is the same as for the first two generations. Our results could not directly highlight implication of the maternal environment on diapause induction, as too few individuals entered diapause in the industrial strain. Yet, maternal thermal conditions did influence development time and fitness-related traits of their offspring, suggesting potential adaptive maternal effects.

### Transgenerational effects on diapause and life-history traits

While we made sure that individuals encountered diapause inducing conditions according to previous studies on *A. ervi* (Christiansen-Weniger and Hardie, 1999; Langer and Hance, 2000; Tougeron et al., 2018b; Tougeron et al., 2018a), our results may highlight insufficiently low temperatures and short periods to reach the thresholds inducing diapause in this industrial strain. Indeed, working on the same industrial strain of *A. ervi* that we used, Mubashir-Saeed *et al.* (in prep.) found 4.7% of non-emerged individuals (that could be dead or in diapause) at the first generation, and an increase to 23.8% at the second, for parasitoids maintained at 8 °C (*vs.* 14 °C for our study) and 8:16 h LD photoperiod. Moreover, at the opposite of wild strains, industrial strains are closed genetic pools maintained under non-diapause inductive rearing conditions over years. This means that these parasitoids are not subject to evolutionary pressures usually maintaining diapause in the natural environment, which may have led to selection on low diapause induction thresholds, as also suggested in wild parasitoid populations from mild winter climates after climate change (Tougeron et al., 2017b).

It could also require more than three generations for recovering diapause in the industrial strain at the tested temperature (14 °C), as shown in two *Trichogramma* species reared under non-diapause-inducing conditions, in which diapause was re-expressed normally (from practically 0% to >50%) after five successive generations under diapause-inducing conditions (Reznik and Samartsev, 2015). Using parasitoids that have undergone diapause to form the next generation could increase the chances of reaching higher diapause levels over time, as it would select lineages forming more diapausing offspring, or with higher sensitivity to diapause-inducing cues. Lengthening of development time over generations for both strains also suggests that parasitoids could show a gradual return of diapause over more generations. This lengthening may involve maternal effects (see below), as the first generation had mothers reared at 20 °C while the following generations had mothers reared at 14 °C. However, as we found some females were never producing diapausing offspring and we observed very low diapause levels (<1%) in the industrial strain, we cannot completely exclude the hypothesis of a genetic loss of diapause in a large part of the population nor the hypothesis of very low diapause induction thresholds in the industrial strain. Knowledge is particularly lacking on the genetic basis of diapause induction thresholds in parasitoids.

We also obtained surprisingly low levels of diapause in the wild strain (a maximum of 21.66 ± 9.97 in the second generation) compared to what was expected for a wild population coming from harsh winter climates (Brodeur and McNeil, 1989; Tougeron et al., 2018a) whereas Tougeron et al., (2018b) found 94.0 ± 2.5 % in similar laboratory conditions (14 °C, 10:14 h LD). This difference in diapause levels of the same population could be explained by the number of generations experiencing standard rearing conditions (20°C, 16:8LD) before being exposed to diapause inducing conditions. Indeed, Tougeron et al., (2018b) tested individuals shortly after they were collected in the field in summer of the same year (September 2015) whereas we maintained this population in constant non-inducing diapause conditions for several generations. A decrease in diapause induction has already been observed after only one (Reznik and Samartsev, 2015) or few (Gariepy et al., 2015; Lauga-Reyrel, 1990) insect generations exposed to non-inducing diapause conditions. However, it is unlikely that diapause has been genetically lost in a large part of the wild population (i.e., that the low diapause levels would be due to batches of females never producing diapausing offspring). Indeed, almost all batches of females produced at least one offspring in diapause, suggesting that in the wild population, high diapause levels could be the result of a maternal response to rearing conditions.

Differences in fitness-related traits of non-diapausing individuals were found between strains and generations. We found that parasitoids from the industrial strain were heavier and contained more fat reserves than the wild strain. Differences between populations may be explained by domestication effect on the industrial strain in which parasitoids could have been selected to increase traits of interest for mass releases (Boivin et al., 2011). Differences between generations may be due to physiological costs of developing and living at 14 °C, which is a sub-optimal temperature for *Aphidius* parasitoids (Sigsgaard, 2000; Le Lann et al., 2011). Decrease in size at the second generation exposed to 14 °C for both strains accompanied by an increase of development time at the second generation do not follow the temperature-size rule in insects, which states that exposure to low-temperature increases adult size (Van der Have and De Jong, 1996). Reduction in adult size has been observed several times in *Aphidius* species experiencing cold stress in both laboratory and natural conditions (Colinet et al., 2007a; Ismail et al., 2012; Tougeron et al., 2016; Wu et al., 2011). This pattern seems to be consistent with the Absolute Energy Demand hypothesis which predicts that larger individuals are disadvantaged under stressful conditions (Blanckenhorn, 2000). Reduction in size was however also observed at 20 °C which indicates a generation effect more than a temperature effect.

### Maternal effect

In our study we could not provide any evidence for maternal effect on diapause induction, mostly due to the very low diapause levels obtained, but we found an increase in development time in the offspring generation, especially when mothers where reared under diapause-inductive conditions (low temperatures and short photoperiod conditions). As discussed before, the industrial strain may have low diapause-induction thresholds that were not reached in this study. In addition, there may be missing signals for maternal effects to be expressed, such as food or humidity (Mousseau and Dingle, 1991; Tauber et al., 1998). However, our results indicated that offspring’s traits and development time were influenced by temperatures experienced by their mothers. Influence of maternal age or temperature and photoperiod experienced by the parental generation on offspring development time have been shown in insects (Mousseau and Dingle, 1991; Fox, 1994). Heritability of development times has also been demonstrated in insects (Sequeira and Mackauer, 1992; Bradshaw et al., 1997; Roff, 2000) and offspring development time can be modified by maternal exposure to different temperatures (Lacour et al., 2014). We observed that the delay in development time (half-emergence) between female parasitoids reared at 14 °C and 20 °C (i.e., 2.7 days) was similar to the delay in development time between their respective offspring, both reared at 14 °C (i.e., 3 days), suggesting an effect of transgenerational plasticity. Morphological and physiological traits were also concerned by these maternal effects, as differences in mean trait value between females from both treatments (14 °C and 20 °C) were passed on their respective offspring. Such maternal effect would prepare offspring to better face incoming environment, as we observed them to be heavier, taller, with wider wings and with more fat reserves.

To conclude, diapause levels expressed in a population and value of life-history traits can be modulated by conditions experienced by precedent generations, and such TGP effects may arise from epigenetic mechanisms or bet-hedging strategies. Natural or artificial selection on plasticity may act on lowering diapause induction thresholds and on modifying the proportion of diapausing offspring produced by each maternal genotype. If the maternal environment is analogous to the offspring’s environment, one can expect that affecting diapause and modifying traits through TGP will result in an adaptive phenotype for the overwintering generation. If not, fitness costs can occur for the offspring, caused by mismatches in the transgenerational transfer of environmental information (mother-offspring conflict, Coleman et al., 2014). In this sense, adaptive significance of maternal effects is still controverted (Marshall and Uller, 2007) because mothers do not always know the best (Henry et al., 2005; Kohandani et al., 2015) and cannot necessarily anticipate incoming environments (Uller et al., 2013). Predicting insect responses to short-term and long-term environmental changes thus implies a better understanding of maternal and transgenerational effects, in particular on fitness-related traits and on phenotypes that are extremely plastic to seasonal variability, such as diapause.

## Acknowledgments

MD performed the experiments. KT and MD wrote the manuscript. All co-authors participated at elaborating the protocol and revising the manuscript. KT was supported by the French Region Bretagne and the Canada Research Chair in biological control. The authors thank technical support provided at the UCL and at the IRBV.

